# Rossmann-toolbox: a deep learning-based protocol for the prediction and design of cofactor specificity in Rossmann-fold proteins

**DOI:** 10.1101/2021.05.05.440912

**Authors:** Kamil Kaminski, Jan Ludwiczak, Maciej Jasinski, Adriana Bukala, Rafal Madaj, Krzysztof Szczepaniak, Stanislaw Dunin-Horkawicz

## Abstract

The Rossmann fold enzymes are involved in essential biochemical pathways such as nucleotide and amino acid metabolism. Their functioning relies on interaction with cofactors, small nucleoside-based compounds specifically recognized by a conserved βαβ motif shared by all Rossmann fold proteins. While Rossmann methyltransferases recognize only a single cofactor type, the S-Adenosylmethionine (SAM), the oxidoreductases, depending on the family, bind nicotinamide (NAD, NADP) or flavin-based (FAD) cofactors. In this study, we show that despite its short length, the βαβ motif unambiguously defines the specificity towards the cofactor. Following this observation, we trained two complementary deep learning models for the prediction of the cofactor specificity based on the sequence and structural features of the βαβ motif. A benchmark on two independent test sets, one containing βαβ motifs bearing no resemblance to those of the training set, and the other comprising 38 experimentally confirmed cases of rational design of the cofactor specificity, revealed the nearly perfect performance of the two methods. The Rossmann-toolbox protocols can be accessed via the webserver at https://lbs.cent.uw.edu.pl/rossmann-toolbox and are available as a Python package at https://github.com/labstructbioinf/rossmann-toolbox.

**Key points:**

- The Rossmann fold encompasses a multitude of diverse enzymes involved in most of the essential cellular pathways
- Proteins belonging to the Rossmann fold co-evolved with their nucleoside-based cofactors and require them for the functioning
- Manipulating the cofactor specificity is an important step in the process of enzyme engineering
- We developed an end-to-end pipeline for the prediction and design of the cofactor specificity of the Rossmann fold proteins
- Owing to the utilization of deep learning approaches the pipeline achieved nearly perfect accuracy

## Introduction

The Rossman fold is one of the most prominent folds in Protein Data Bank and by far the most functionally diverse one, with >300 different functions, typically involving the addition of a methyl group on a substrate (methyltransferases) or transfer of electrons from one molecule to another (oxidoreductases) [1–3]. It is also assumed to be one of the oldest folds, which was already well represented in the last universal common ancestor (LUCA). From the structural perspective, the Rossmann fold belongs to the general class of β/α proteins and comprises four connecting α-helices and six consecutive β-strands (arranged in the 3-2-1-4-5-6 order) forming a parallel pleated sheet (Figure 1). Rossmann-fold enzyme families are characterized by their use of cofactors, and in particular of nucleoside-containing cofactors such as S-Adenosylmethionine (SAM), nicotinamide adenine dinucleotide (NAD), nicotinamide adenine dinucleotide phosphate (NADP), flavin adenine dinucleotide (FAD), and others. These cofactors share not only the biochemical compound (adenosine) but also bind to the same specific region of the Rossmann fold, even in distantly related proteins. The cofactor-binding site shared by all members of the Rossmann fold corresponds to a small structural fragment comprising β1–α1–β2 and the connecting loops [4]. Interestingly, this fragment has been identified as one of the ancestral peptides [5] that may have existed as a nucleotide-binding unit even in the pre-LUCA times. However, beyond the shared homologous cofactor-binding motif, many of the Rossmann enzymes do not show detectable homology all along the sequence, and the greater part of their sequences has diverged beyond recognition.

**Figure 1.**
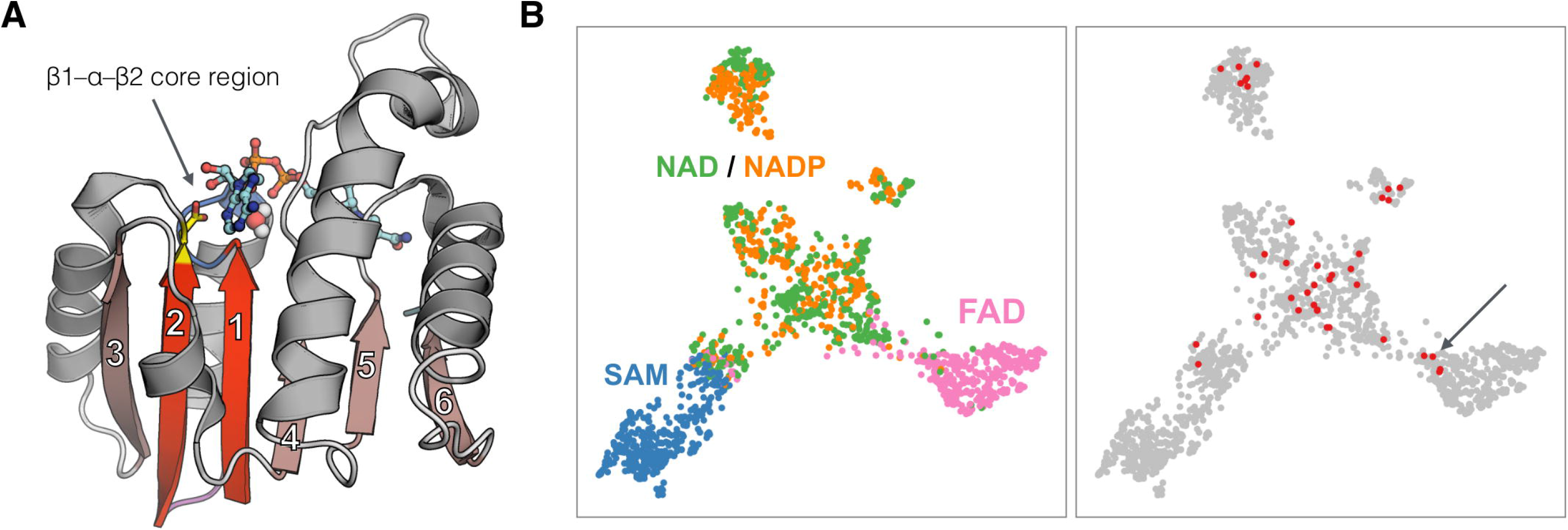
Cofactor recognition in proteins adopting the Rossmann fold. **(A)** Example of a Rossmann fold protein, the malate dehydrogenase from *Escherichia coli* bound to NAD cofactor (shown as a ball-and-stick model). Beta strands are numbered according to the topological order and the two of them that participate in the formation of the βαβ cofactor-binding motif are indicated with a brighter color. The aspartic acid residue essential for the cofactor binding is shown in yellow. **(B)** Sequence-based clustering of Rossmann βαβ motifs used to train and test the two prediction models. Points correspond to 1,647 βαβ motifs and their positions reflect the relative sequence similarity. The left panel depicts core regions colored according to the bound cofactor type, whereas the right panel highlights core regions (shown in red) used for benchmarks based on experimental data.

The NAD, NADP, and FAD cofactors are essential for the functioning of oxidoreductases, whose role is to transfer electrons from one molecule (electron donor) to another (electron acceptor). For example, alcohol dehydrogenases facilitate the oxidation of alcohol (electron donor) to aldehyde with the concurrent reduction of NAD+ (electron acceptor) to NADH. Generally, NAD occurs mostly in catabolic reactions, i.e., reactions that lead to the decay of complex molecules, and as a result, produce energy, whereas NADP (differing from NAD only by an additional phosphate group) is involved mostly in anabolic reactions, which create complex molecules from simple substrates and thus store energy. The addition of the phosphate in NADP does not alter its electron transport capability; however, the phosphate group modifies the structure of the cofactor, which allows different enzymes to have different specificities for NAD and NADP, thereby decoupling the catabolic and anabolic reactions [6]. In contrast to NAD(P) and FAD, SAM takes part in methylation reactions, i.e., transferring a methyl group from SAM to substrates like DNA/RNA, proteins, or small-molecule secondary metabolites [7], in turn, its derivative decarboxylated SAM (dcSAM) plays a role in pathways of the polyamine biosynthesis [8].

The rational cofactor specificity re-engineering is used for manipulating metabolic pathways [9,10], and it has applications in drug engineering and industry [6]. One of the first attempts to redesign the cofactor specificity of a Rossmann-like enzyme was a work by Scrutton and colleagues [11]. By investigating the *Escherichia coli* glutathione reductase, the authors identified amino acids that confer specificity for NADP and then systematically replaced them to achieve cofactor preference gradually switched towards NAD, while preserving the specificity towards the substrate. To this date, there were many other successful attempts to rationally change the cofactor specificity of Rossmann enzymes [12]; however, most of them were based on experimental or theoretical structures of the target protein and/or a detailed sequence alignment among the family members [13–15]. These successful cases of NAD to NADP and *vice versa* conversions were the basis for the formulation of rules defining how properties of amino acids located at the cofactor-recognizing site dictate its binding specificity [16].

The extensive research on the cofactor specificity determinants has led to the development of universal computational models. For example, Cui et al. proposed an approach in which molecular dynamics simulations were used to evaluate mutants based on their propensity to form hydrogen interactions with a cofactor [17]. A structure-based strategy was also employed in CSR-SALAD, a method that aids the selection of amino-acid positions for the site-saturation mutagenesis [18], and in MaSIF, a recently developed deep learning framework for the identification of structural fingerprints important for protein-ligand interactions [19]. Cofactory is the only available computational tool capable of high-throughput, sequence-based evaluation of Rossmann enzymes for their ability to bind NAD, NADP, and FAD cofactors [20]. However, the method does not consider SAM, and its accuracy is far from satisfactory, especially in the case of NADP-preferring proteins.

Obtaining accurate predictions for a wild-type sequence and its potential variants is a prerequisite for cofactor re-engineering tasks. However, performing such analyses with the currently available approaches requires a time-consuming, case-by-case investigation of relevant sequences, structures, and literature. To address this problem, we collected all known experimental structures of the Rossmann fold proteins complexed with cofactors and used this data to train deep-learning-based models for the prediction of the cofactor specificity in Rossmann enzymes based on the sequence or structure of the βαβ motif. We rigorously tested the methods using a test dataset comprising examples sharing no more than 30% sequence identity to the dataset used for the training and a panel of 38 experimentally confirmed transitions between NAD and NADP enzymes. Both benchmarks revealed very good accuracy of the models and their applicability to re-design tasks.

## Methods

### Data preparation

From 44 representative Rossmann structures selected based on the literature [4] and the ECOD [21] classification (Supplementary Table 1), we extracted C◻ atoms corresponding to *3-2-1-4-5* β-sheets and the α-helix connecting β1 and β2 (Figure 1). These partial backbone structures were used to search the Protein Data Bank using MASTER [22]. The resulting matches were processed using Python scripts to obtain fragments corresponding to the βαβ motifs responsible for the cofactor binding. For handling the structures, we used Atomium [23] and *localpdb* (Ludwiczak et al., https://github.com/labstructbioinf/localpdb, manuscript in preparation). The motifs were analyzed with PLIP [24] to identify protein-cofactor interactions, and all the motifs lacking such interactions were discarded. Finally, we defined labels for use in machine learning by combining cofactor variants: NAI, NAJ, NAD to NAD; NAP and NDP to NADP; FAD, FDA to FAD; and SAM, SAH to SAM.

The resulting set of 11,487 redundant cofactor-bound βαβ motifs was clustered with mmseqs2 [25] (min. sequence identity 0.3, coverage 0.5, coverage mode 1, clustering mode 2), yielding 483 clusters comprising 1,647 unique βαβ motifs (Figure 1). The training, validation, and test sets contained 68%, 16%, and 16% of these βαβ motifs, respectively. We maintained a balance within the training set (the approximately equal number of examples for each cofactor class), and between the validation and test sets (to make sure that the performance estimated on the validation set is a good approximation of the performance on the test set). Most importantly, we ensured that the train, test, and validation sets are separate in the sense that maximal sequence identity between any pair of βαβ motifs originating from two different sets never exceeds 30%. The detailed statistics of the individual sets can be found in Supplementary Table 2.

### Prediction models

We considered two complementary approaches to tackle the problem of cofactor specificity prediction in Rossmann fold proteins. Both rely on deep learning procedures but differ in terms of the neural network architectures and data type used. The first uses only sequences of βαβ motifs, whereas the second employs also the structural data represented in the form of graphs (Figure 2). The training and validation sets were used to train the models and select the optimal ones, respectively, whereas the test set (comprising βαβ motifs showing no more than 30% sequence identity to the βαβ motifs from other sets) was used for estimating the effectiveness of ours and previously developed [19,20] models. In the following sections, we provide a basic description of the structure and sequence-based models. For more details, please refer to the Supplementary Methods.

**Figure 2.**
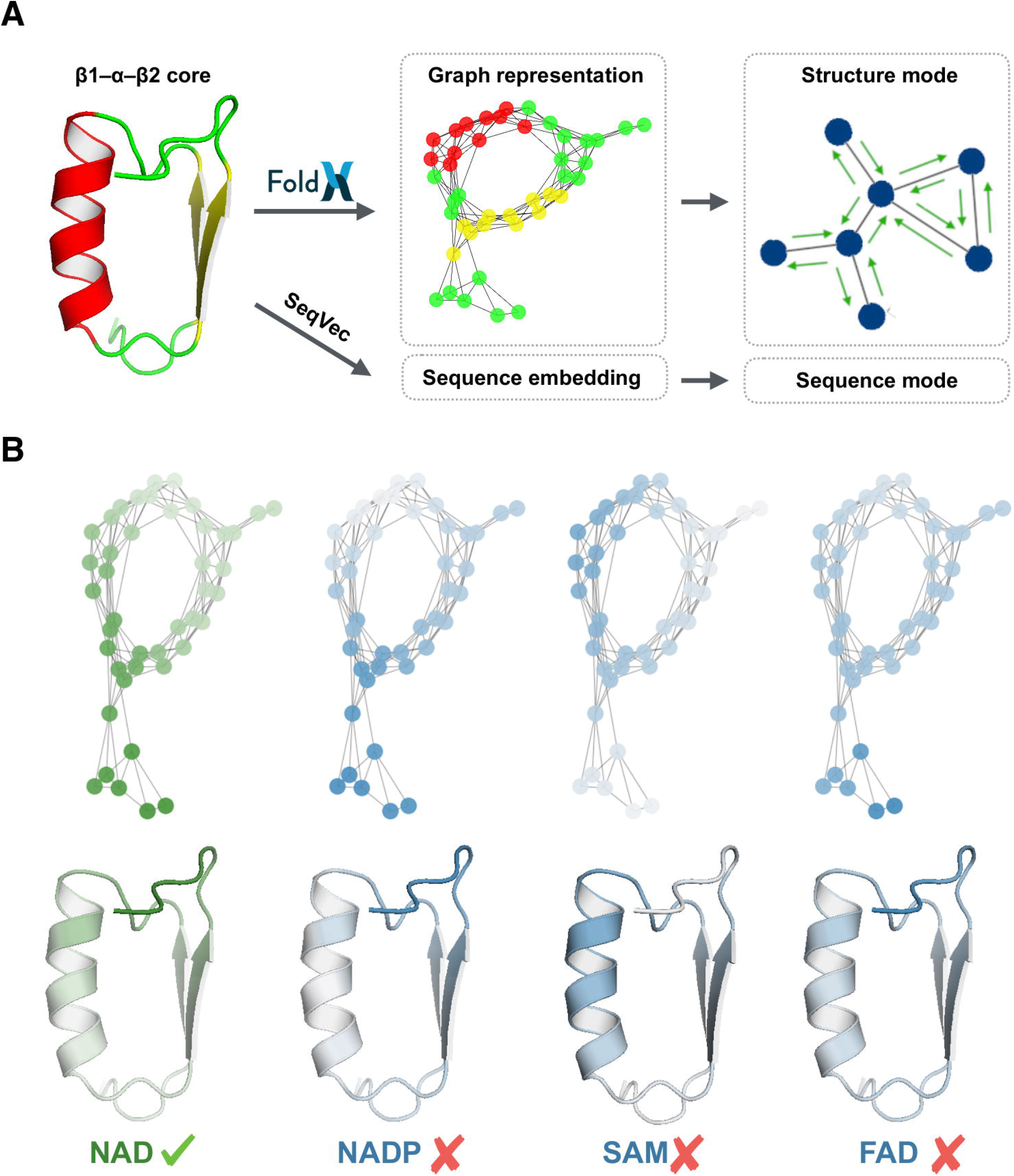
General scheme of the prediction pipeline. **(A)** The pipeline consists of two prediction models, which enable the cofactor specificity prediction based on the sequence or structure of the βαβ core. **(B)** The prediction models return not only the binding probabilities but also per-residue importance scores reflecting the individual residues’ contribution to the final prediction. Colors ranging from green via white to blue indicate positive, neutral, and negative impact on a given prediction, respectively.

To develop the sequence-based predictor, all the sequences from the train, validation, and test sets were embedded with the SeqVec [26], resulting in the vectors of size [N, 1024] (where N is the length of the βαβ motif sequence). Neural network architecture was adopted from the original SeqVec paper [26], which described several applications of the embeddings for sequence classification tasks. Briefly, the SeqVec embeddings were processed by two consecutive convolutional layers and connected through two densely connected layers to the sigmoid-activated 4-class output layer denoting the binding probability for each of the cofactor classes. Batch normalization and random dropout (probability 0.5) operations were applied after each convolutional layer to avoid overfitting. The training was performed for 50 epochs with the cross-entropy loss function, one-hot encoded labels derived from the structural data (see the preceding section for the details), and the Adam optimizer [27] as implemented in the *tensorflow* Python package. Input vectors were centered and zero-padded to the constant length of 65. Model weights were saved from the epochs corresponding to the highest macro-F1 score on the validation set. The top 10 models, exhibiting the highest macro-F1 scores on the validation set, were used to create the final ensemble, which averages the outputs of these best-performing models. The per-residue contributions to the predicted cofactor binding classes were calculated using the *captum* Python package with the *integrated gradients* method [28] implemented therein.

The structure-based predictor was developed using graph neural networks enabling a more natural representation of complex non-grid data, such as protein structures (Figure 2). An undirected graph G is defined as a set of nodes N, also termed vertices, n ∈ N, and a set of edges E; if two vertices are connected by an edge then *e*_*ij*_ = *e*_*ji*_ ∈ *E*. The βαβ motif dataset structures were converted to graphs in which nodes represented the individual residues and edges defined interactions between them (two residues were considered to be interacting, i.e., forming an edge, when the distance between their C◻ atoms was below 7 ◻). Subsequently, nodes and edges of the graphs were annotated with structural data. To this end, the structures containing the βαβ motifs were minimized in the FoldX force field [29] using the *RepairPDB* command, then the structural features were extracted with *SequenceDetail* and *PrintNetworks* commands and assigned to nodes and edges, respectively (Supplementary Table 3). Such a graph representation constituted the input to the network, whereas its output was a four-element vector reflecting the probability toward binding of the individual cofactors.

The graph neural network model was implemented in Deep Graph Library [30] using PyTorch backend and Lightning training routines. It is composed of a series of EdgeGAT layer blocks, where each block contains an EdgeGAT layer (see Supplementary Methods for details) followed by a batch normalization layer (edges and nodes are treated separately) with a LeakyReLU activation function. To transform graphs of various sizes to a fixed-size representation, the last EdgeGAT layer produces the output with node features of size 4 (i.e., number of cofactor classes), which are subsequently summed over the graph and passed through the fully connected layer followed by the Sigmoid activation function. The two main hyperparameters of the network are the number of internal EdgeGAT blocks and the size of the node features vector (the edge features size was fixed at 20). For the training, the focal loss cost function [31] and Adam optimizer [27] with L2 regularization were used. Training stopping criteria were given by not increasing the F1-macro score over the validation set. In total, we trained 1,400 models with various hyperparameter values (two to four EdgeGAT blocks and the size of node features ranging from 32 to 512), and the four models that performed the best on the validation set were selected to build the final ensemble model.

### Benchmark of the prediction models

We benchmarked MaSIF-ligand [19], Cofactory [20], and the two methods developed in this study. Sequences were used as inputs for Cofactory and the sequence-based predictor, whereas the corresponding structures were used for the MaSIF-ligand and the structure-based predictor. The performance was assessed using two data sets: the test set (see “*Data preparation*” above) and an auxiliary test set built based on 38 published mutagenesis studies aiming at a change of the cofactor specificity from NAD to NADP and *vice versa* (Supplementary Table 4). For the benchmarking of MaSIF-ligand and Cofactory, we ensured that the entries (structures and sequences, respectively) used to train these methods were excluded. Moreover, to verify whether the auxiliary test set is not biased, we performed clustering of all its sequences together with the sequences from the train-test-validation set. To this end, we calculated a SeqVec [26] embedding for each βαβ motif sequence, compared them all-vs-all using cosine similarity metric, and used the resulting matrix as an input to the UMAP [32] dimensionality reduction procedure (Figure 1).

### Per-residue contribution scores

The models developed in this study predict the cofactor binding specificity and return scores indicating the contribution of the individual residues to the prediction (Figure 2). To use such data in guiding the protocols for the prediction of the specificity-switching mutations (see below), we used the following procedure. First, the initial set of 11,487 redundant βαβ motifs bound to cofactors (see “*Data preparation*” for details) was subdivided into four groups corresponding to the four cofactors. Then, in each group, the redundancy was removed with mmseqs2 [25] (min. sequence identity 0.7, coverage 0.5, coverage mode 0, clustering mode 0) and a structure-based multiple sequence alignment (MSA) was calculated with parMATT [33]. Finally, all the constituent βαβ motifs were evaluated with the structure and sequence-based prediction models, and the average sequence and structure-based importance scores for each column in each MSA were calculated. In this way, by aligning a βαβ motif sequence (WT or mutated) to the MSA associated with a given cofactor, one can infer which residues would be (or are) crucial for its recognition.

### Brute-force mutational scan protocol

For each wild-type βαβ motif of the auxiliary test set (see “*Benchmark of the prediction models*” above), all possible point mutations were defined and the resulting variants of full-length Rossmann domains were modeled with Modeller [34] and FoldX [29]. Subsequently, the affinity of the mutants towards NAD and NADP cofactors was predicted using the sequence and structure-based models. The most plausible mutants, i.e., those which may change the cofactor specificity in the assumed direction, were selected using the following procedure. First, possibly unstable models characterized by FoldX ddG score or Modeller DOPE score greater than 4.5 and 240, respectively, were discarded. Then, the raw scores for NAD and NADP were adjusted by multiplying them by the corresponding importance scores associated with the position where a given mutation was introduced. Finally, for each case, the mutants were sorted according to the adjusted scores, and the positions of experimentally confirmed mutants were indicated.

### Iterative mutational scan protocol

For the prediction of the specificity-switching variants involving more than one mutation, an iterative mutational scan protocol was developed. In contrast to the brute-force protocol in which all possible variants are evaluated exhaustively, the iterative protocol relies on Monte Carlo simulations during which residues to be changed are selected according to the contribution scores obtained for the target cofactor. A detailed description of the procedure can be found in the Supplementary Methods.

The iterative mutational scan protocol was evaluated using the auxiliary test set. To this end, for each benchmark case, sequences from all Monte Carlo simulation steps were binned based on the actual number of mutations relative to the WT. In each group, the 95th percentile of the score was defined and variants with scores below this value were discarded. The remaining sequences from all bins were collected and the 20 most frequent point mutations were determined (in the case of sequences containing more than one mutation, each was treated separately). Such a list was then filtered by removing variants with FoldX ddG score above 4.5 or Modeller DOPE score above 240, and sorted by the frequency of the individual mutations. An analogous procedure was used to determine the most frequently occurring pairs of mutations. In this case, all the variants that remained after FoldX and Modeller filtering were analyzed to determine co-occurring mutations (in the case of variants containing more than two mutations, all possible combinations were considered regardless of the relative position in the sequence). Then, the pairs not involving the mutations from the previously defined top 20 list were removed and the remaining ones were sorted according to their frequency, resulting in a ranking of mutations’ co-occurrence.

## Results and Discussion

### The minimal cofactor specificity-defining region

The most conserved interactions between the Rossmann fold proteins and their cofactors occur in the region corresponding to the βαβ motif [4]. Consequently, mutating the residues in this region is typically sufficient to alter the cofactor specificity (Supplementary Table 4). To gain insight into the determinants of cofactor specificity, we performed a clustering analysis of βαβ motif sequences (Figure 1). We found that most of the SAM and FAD-binding motifs were contained in distinct and separate clusters (blue and magenta, respectively), whereas the NAD and NADP-binding motifs (green and orange, respectively) were mixed in the central supercluster. This suggests the existence of specific features of the SAM and FAD-binding motifs and confirms that the evolutionary transition between NAD and NADP binding occurred multiple times.

Regardless of whether the specificity-determining features are of divergent or convergent origin, we see challenges in employing them for the development of tools for the design and prediction of cofactor specificity. First, considering that even a few point mutations are capable of transforming the specificity of NAD and NADP-binding motifs, distinguishing them requires the reside-level resolution. Second, despite their apparent distinctiveness, some of the SAM and FAD-binding motifs show significant similarity to NAD(P) binders, making their proper annotation difficult. Finally, it is not clear whether the sequence features alone are sufficient, or the usage of additional structural data would be beneficial.

### Predicting the cofactor binding specificity

Given the above considerations, we reached for modern deep learning techniques and developed two models for the prediction of the cofactor binding specificity based on either the sequence or the structure of βαβ motifs (see Methods). For a given βαβ motif, the models predict the binding probability for each cofactor and importance scores defining which residues were essential for the predictions (Figure 2). To assess the performance of the methods, we built a separate test set comprising βαβ motifs with known binding specificity and sharing no more than 30% sequence identity to those used to train the methods. In such a benchmark, the sequence and structure-based predictors achieved very good accuracies (93% and 94%, respectively; Figure 3) and outperformed the currently available methods, i.e., Cofactory (61%) [20] and MaSIF-ligand (58%) [19]. While the poor performance of Cofactory was to be expected, as the method was developed more than 15 years ago, the weak predictive power of a MaSIF-ligand was surprising. Close inspection of the results revealed that in ~40% of the test set examples, the MaSIF-ligand crashed during the execution of the external tools such as those for the calculation of surface electrostatics. To account for this effect, we also evaluated the method using only the test set entries that caused no problems and found that it achieved a macro-F1 score of 85%. The fact that even after removal of the problematic cases, MaSIF-ligand did not achieve performance comparable to the methods developed in this study may be partially related to the data preparation procedure. The MaSIF-ligand was only trained on NAD, NAP, FAD, and SAM ligands, whereas our pipeline captures also their variants (see “*Data preparation”* section for details). This result also indicates that the performance of bioinformatics tools can be considerably hampered by technical issues. Bearing this in mind, we limited to the minimum the external dependencies of our tools and developed a web server, providing easy access for non-expert users (https://lbs.cent.uw.edu.pl/rossmann-toolbox).

**Figure 3.**
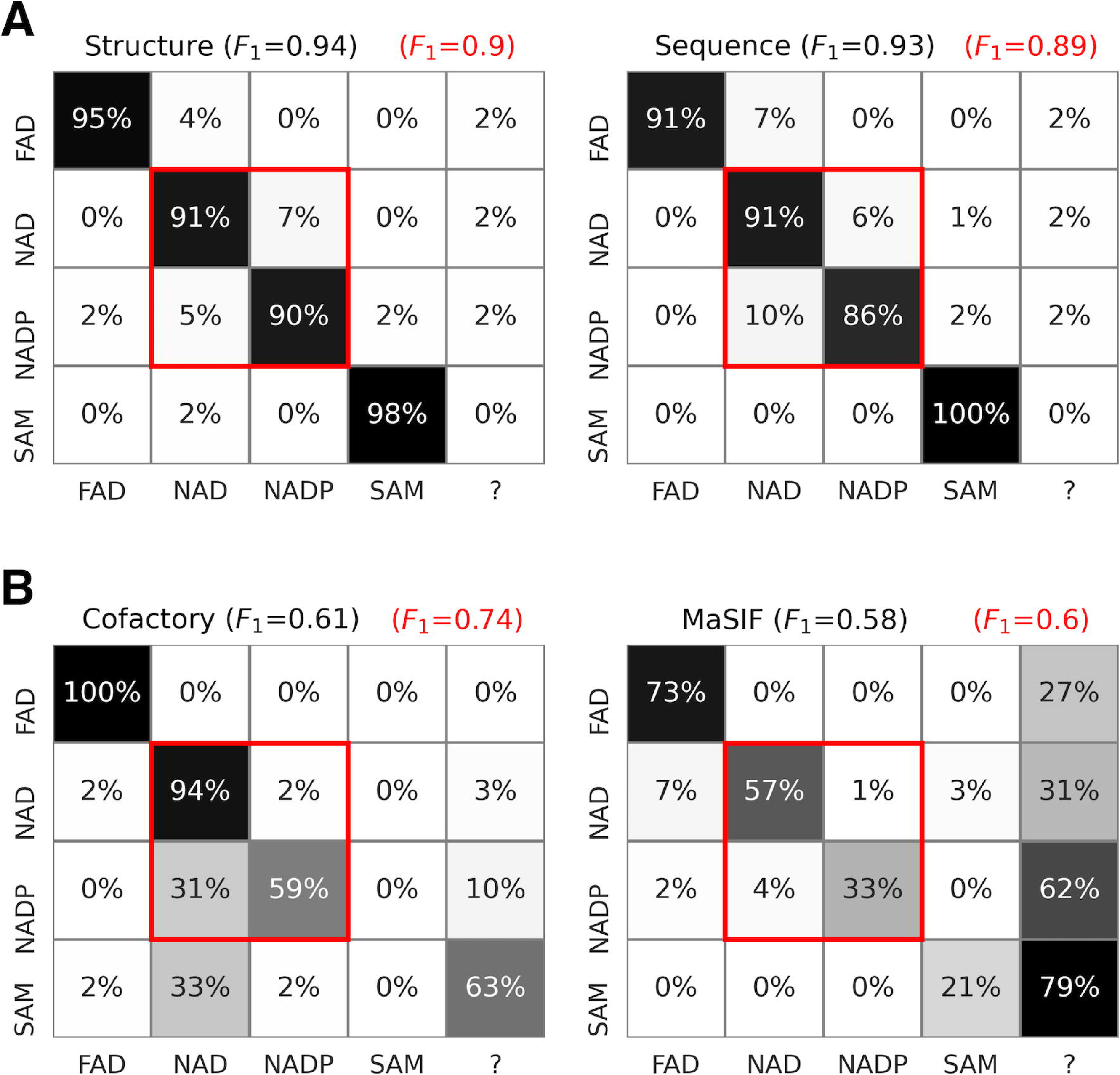
Evaluation of the prediction models using an independent test set. Each confusion matrix corresponds to a single method. The vertical and horizontal axes denote ground truth and predicted binding, respectively. The category indicated with a question mark defines cases for which no prediction could be obtained. **(A)** Performance of sequence and structure-based models developed in this study. **(B)** Performance of other available tools, MaSIF-ligand and Cofactory.

### Predicting the effects of mutations

Despite the good performance on the test set, we deemed it necessary to validate our methods on more difficult, real-life examples. To this end, we built an auxiliary test set comprising 38 published experiments that aimed at switching the cofactor specificity from NAD to NADP or *vice versa* by introducing one or more mutations in the βαβ region (see Methods). Importantly, these cases are representative (red points in Figure 1) and exemplify a variety of mutation design approaches ranging from loop exchange [35,36], evolutionary-based [37] to computational predictions [17,18].

For each βαβ motif pair (WT and mutated) of the auxiliary test set, we calculated ΔNAD and ΔNADP scores quantifying the mutation-induced changes in the predicted binding probabilities. Examination of these scores revealed a perfect performance of our methods, both of which achieved 100% accuracy in predicting the direction of specificity change (Figure 4). Moreover, we checked the accuracy of the methods in predicting the preferred cofactor of the WT βαβ motifs and found that the structure and sequence-based methods performed well (92% and 97%, respectively; Figure 4). For this benchmark, we did not consider mutated sequences because the increase in the affinity towards one cofactor does not necessarily imply a decrease in the affinity towards the other, and the resulting mutated enzymes may have dual specificity, e.g., [14,17]).

**Figure 4.**
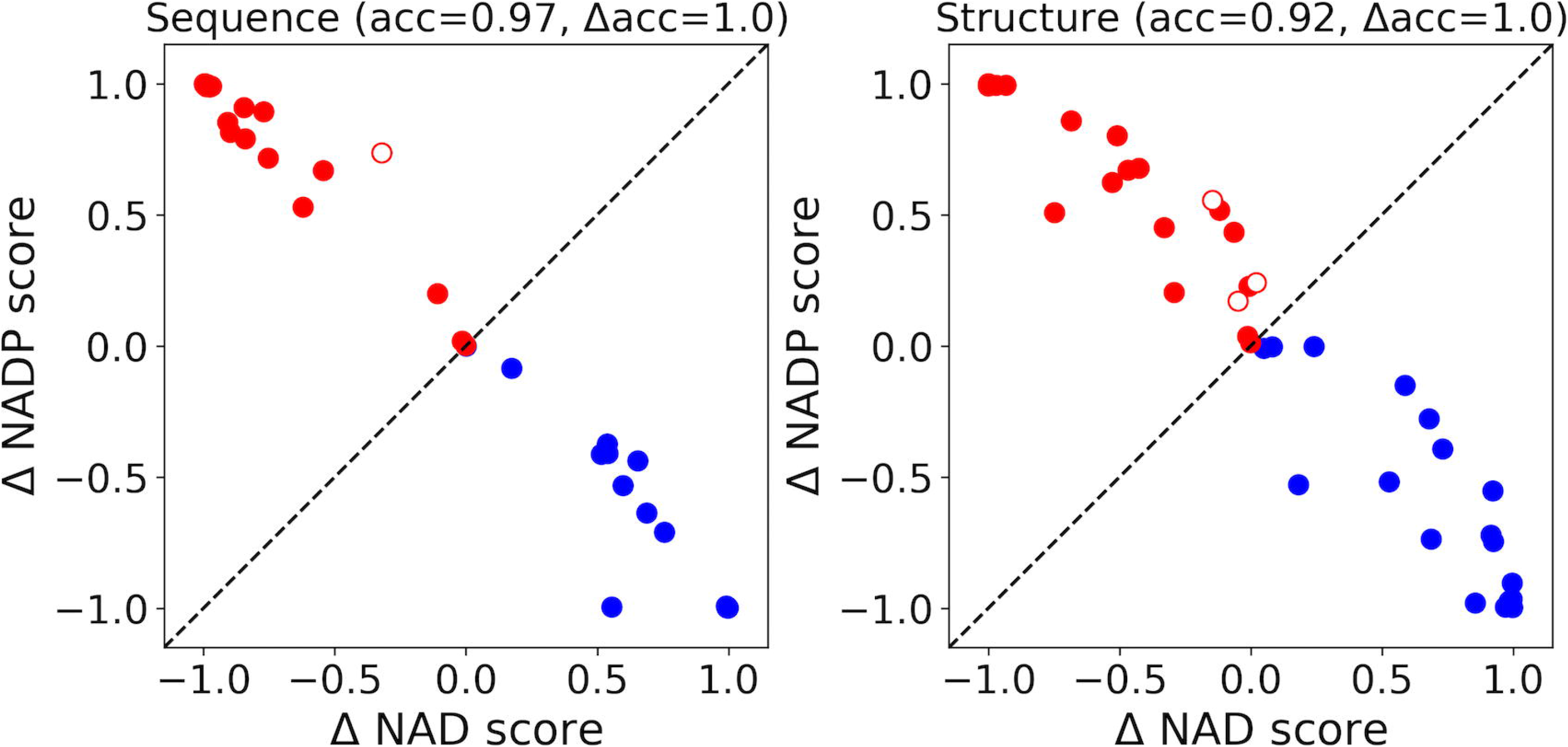
Evaluation of the prediction models developed in this study using the auxiliary test set. Circles correspond to experimentally confirmed cases of changing the cofactor specificity between NAD and NADP by one or more mutations. They are colored according to the direction of change: NAD to NADP and NADP to NAD are indicated with red and blue colors, respectively. Empty circles denote cases for which the change of the specificity was predicted correctly but the specificity of the WT was mispredicted (see text for details). ΔNAD and ΔNADP denote the difference between predicted binding probabilities of WT and mutated variants.

We identified only three cases in which one or both of our methods failed to predict the cofactor specificity of the WT βαβ motif. The first one, dihydrolipoamide dehydrogenase, an E3 component of the pyruvate dehydrogenase complex [38], contains two Rossmann fold domains, both belonging to the FAD/NAD(P)-binding group defined in the ECOD database [21]. The first domain is involved in FAD binding, whereas the second recognizes NAD. The proposed mutations [39] aimed at switching the cofactor specificity of the latter domain to NADP. The sequence and structure-based models correctly predicted the effect of these mutations; however, both predicted the WT βαβ motif to bind FAD instead of NAD. The second mispredicted case, water-forming *Streptococcus mutans* NADH oxidase, has the same domain composition as the dihydrolipoamide dehydrogenase and contains two Rossmann domains from the FAD/NAD(P)-binding group. Also, in this case, the second Rossmann domain was mutated [40] to change its specificity from NAD to NADP and the effect of these mutations was predicted correctly but the WT βαβ motif obtained the highest score for FAD instead of NAD (in this case, however, only the structure-based method yielded such a result). The fact that in both cases the FAD score exceeded the NAD score can be attributed to the evolutionary position of the respective βαβ motifs. Both, despite being experimentally confirmed NAD binders, cluster together with FAD-bound βαβ motifs (arrow in Figure 1) and thus may constitute an example of the specificity switch within the FAD/NAD(P)-binding group. Since such cases are rare, they may have been missed during the training process. The last mispredicted case, the flavoprotein monooxygenase, can utilize both NAD and NADP [41], thus it is not surprising that the predictions were ambiguous.

### Predicting the cofactor binding affinity

Although the accuracies of the sequence- and structure-based models turned out to be nearly identical (Figures 3 and 4), their predictions differed, especially in the most uncertain cases (Supplementary Figure 1). Such a partial lack of correlation indicates that, to some extent, the methods could have captured different aspects of the cofactor specificity determinants. While examining this, we noted that the structure-based predictions for specificity-switching variants of the auxiliary test set tend to correlate with their corresponding kinetic constants expressed in terms of the k_cat_ to K_m_ ratio. For example, Scrutton and coworkers designed 10 variants of the βαβ motif of *Escherichia coli* glutathione reductase that gradually changed the specificity of the enzyme from NADP to NAD [11]. This transformation was reflected in the structure-based predictions, which not only distinguished the extreme variants but also the intermediate ones (Figure 5). An analogous pattern was observed in the predictions for *Candida boidinii* formate dehydrogenase variants aiming at switching the specificity from NAD to NADP. Also, in this case, the gradual switch towards the target cofactor could be seen both in the experimental data and *in silico* predictions (Figure 5). These results indicate that the utilization of additional structural data was beneficial as it resulted in a more versatile (yet slower) prediction model.

**Figure 5.**
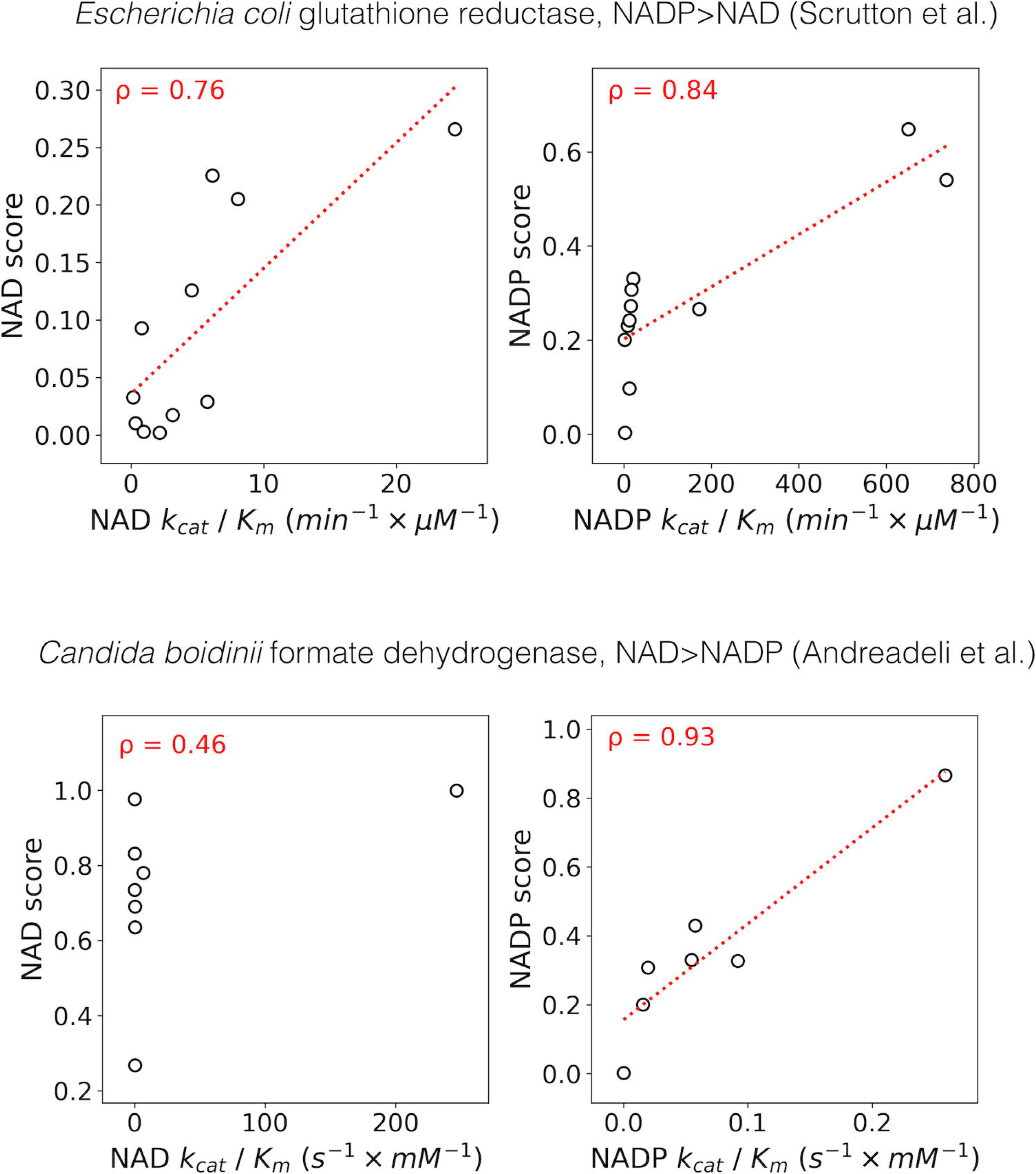
The relationship between the structure-based predictions and kinetic constants expressed in terms of the k_cat_ to K_m_ ratio. In each panel, the vertical axis denotes the predicted binding score for a given cofactor, whereas the horizontal axis depicts the corresponding experimental values.

### Designing the specificity-switching mutations

In the benchmarks described above, we estimated the ability of our models to predict the cofactor specificity and its change upon mutation. In such cases, however, both the wild-type and mutant sequences are known. To mimic real-life scenarios in which the cofactor-switching mutations for a given βαβ motif are predicted from scratch, we reached for two approaches: one relying on the evaluation of all possible point mutations (brute-force approach) and the other employing Monte Carlo heuristic to identify complex variants in which more than one position is altered (iterative approach); see Methods for details.

To test the brute-force approach, each WT sequence of the auxiliary test set was used as a starting point to generate all possible point mutations (19\**n*, where *n* is the length of the βαβ motif). Then, the mutations were separately evaluated with the structure and sequence-based models, sorted according to the predicted affinity towards the target cofactor, and the position of the correct mutation, i.e., the experimentally confirmed one, was indicated (Figure 6). The lower is the position of the correct mutation in the ranking, the fewer lab experiments would have been necessary to reveal it, and the more effective is a given method in *de novo* prediction of cofactor specificity-switching mutations. For example, for eight out of nine cases, in which single substitutions were sufficient to change the specificity (“*p*” in Figure 6), both models identified the correct mutations among 1% of the top-scored variants. Similarly, for most “complex” cases featuring two or more mutations (“*c*” in Figure 6), at least one of them was identified among the 1% top-scored.

**Figure 6.**
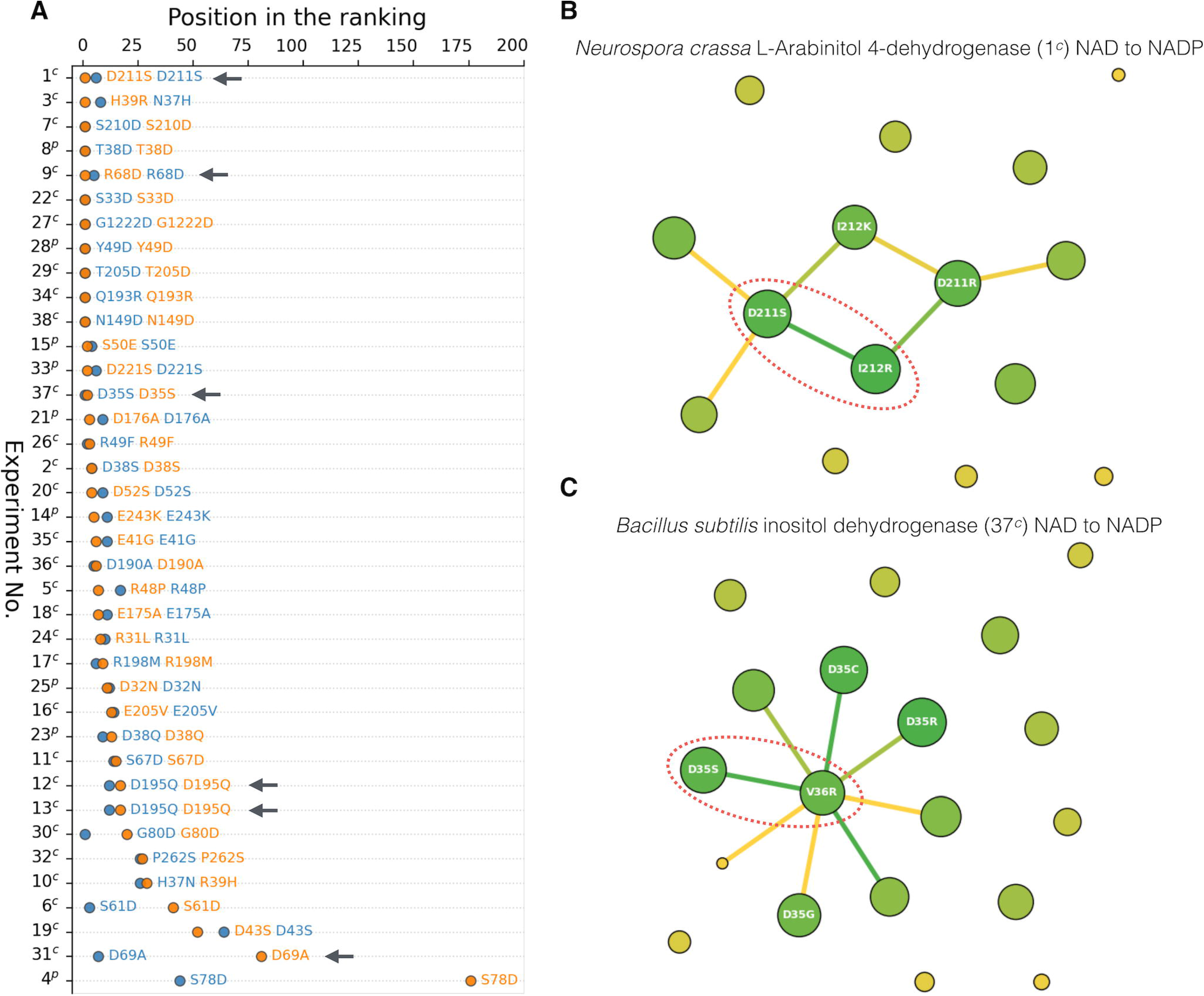
Performance of the sequence and structure-based models in the task of cofactor specificity design. **(A)** Results of the brute-force mutational scan of the 38 βαβ motifs from the auxiliary test set. Rows correspond to the 38 experiments in which the specificity change was achieved by either point or complex (double, triple, etc.) mutations (“*p*” and “*c*” suffixes, respectively). The orange and blue circles (sequence and structure-based model, respectively) indicate the position of the experimentally confirmed mutations in respect to the ranking of all possible point mutations ordered according to the predicted affinity towards the target cofactor (the lower the position, the better performance). In experiments relying on complex mutations, only the first best-scored mutation is shown. **(B)** Result of the iterative mutational scan of L-Arabinitol 4-dehydrogenase. Circles denote mutations (their sizes are proportional to the frequency of occurrence), whereas edges between them define predicted coupling (the greener is an edge, the more probable is the given coupling). Experimentally confirmed pairs of mutations are indicated with red ovals. **(C)** Result of the iterative mutational scan of inositol dehydrogenase.

Among the 29 “complex” cases, six relied on a multi-step approach in which mutations were iteratively added and tested experimentally to obtain increasing specificity towards the target cofactor (1^*c*^, 9^*c*^, 12^*c*^, 13^*c*^, 31^*c*^, and 37^*c*^; indicated with arrows in Figure 6). For example, the study aiming at switching the cofactor specificity of L-Arabinitol 4-dehydrogenase (1^*c*^) from NAD to NADP [42] involved multiple rounds of rational design. The D211S variant obtained at the first round showed a decrease in activity towards NAD, with a minimal yet detectable activity increase towards NADP, whereas the second-round double mutant D211S/I212R displayed actual reversal in cofactor specificity. The brute-scan approach identified the first-round D211S variant with the highest confidence. Intrigued by this observation, we investigated the remaining cases and found that in all but one of them (9^*c*^) the first-round mutation was also the one with the highest rank, demonstrating the ability of the brute-force scan to identify the key mutations.

Using the brute-force approach for the identification of all mutations in the “complex” cases would be computationally infeasible. To address this problem, we have developed an iterative approach capable of simultaneous prediction of multiple mutations. In contrast to the brute-force scan, this procedure returns not only a ranking of specificity-switching mutations but also of their co-occurrences. For example, in the case of the aforementioned L-Arabinitol 4-dehydrogenase (1^*c*^), the D211S mutation indicated by the brute-force scan was predicted to co-occur with two mutations I212K and I212R (Figure 6), and the one showing the strongest coupling (I212R) was also confirmed experimentally [42]. Another example was the reengineering of the cofactor specificity of *Bacillus subtilis* inositol dehydrogenase (37^*c*^) from NAD to NADP (Figure 6). In this case, our predictor identified the coupling between two out of three mutations (D35S and V36R) that were revealed by the experimental study [15]. The third mutation, A12K, was not predicted; however, it must be noted that the double mutant D35S/V36R already preferred NADP over NAD by a factor of 5. In fact, the A12K mutation was not essential, and its purpose was to improve the specificity change further. Inspection of all 29 benchmark cases featuring two or more mutations revealed that in 15 of them the iterative approach predicted more than one correct mutation, and in 8 out of these 15 cases also the correct coupling between them (Supplementary Table 5).

### Conclusions

Sequence embeddings and graph neural networks are relatively new tools, and their applicability to biological tasks is being explored. In this study, we demonstrated their usefulness in the accurate prediction of protein-cofactor interactions in Rossmann fold proteins and developed new solutions for the non-lossy representation of protein structures for machine learning purposes (see Methods and Supplementary Methods for details). The usage of additional structural data did not drastically improve the accuracy (Figure 3), however, it enabled predictions of the relative binding affinity (Figure 5). We envision that for not yet attempted reengineering tasks, such as a switch between NAD(P) and SAM, it may be necessary to use structural descriptors capable of capturing subtle features [43] otherwise disregarded by the sequence-based method. Finally, it is important to note that our methods were trained only with natural βαβ motifs, which makes them prone to assign spurious scores to non-natural variants that are wrong from the structural perspective. This problem was overcome by utilizing Modeller and FoldX energy functions to detect and discard potentially unstable variants. However, a more elegant solution would be introducing such variants to the training set and marking them as non-binders. This and other issues will be addressed in course of the development of subsequent releases of the Rossmann toolbox.

## Authors’ contributions

SDH, KK, and JL designed the study. SDH, KK, JL, RM, and KS prepared the datasets. KK designed and implemented the graph-based prediction model, whereas JL designed and implemented the sequence-based prediction model. MJ and AB designed and implemented the MC-based mutational scan. KS implemented the webserver. SDH, KK, JL, and MJ drafted the manuscript. All authors read and approved the final version of the manuscript.

## Acknowledgments

This work was supported by the First TEAM program of the Foundation for Polish Science co-financed by the European Union under the European Regional Development Fund (grant POIR.04.04.00-00-5CF1/18-00 to SDH). JL was in part funded by the National Science Centre (grant 2017/27/N/NZ1/00716 to JL). The authors would like to thank Paola Laurino, Saacnicteh Toledo Patino, and Vikram Alva for their constructive comments and critical evaluation of the manuscript.

## Bibliography

1. Tóth-Petróczy A, Tawfik DS. The robustness and innovability of protein folds. Curr. Opin. Struct. Biol. 2014; 26:131–8

2. Medvedev KE, Kinch LN, Schaeffer RD, et al. Functional analysis of Rossmann-like domains reveals convergent evolution of topology and reaction pathways. PLoS Comput. Biol. 2019; 15:e1007569

3. Medvedev KE, Kinch LN, Dustin Schaeffer R, et al. A Fifth of the Protein World: Rossmann-like Proteins as an Evolutionarily Successful Structural unit. J. Mol. Biol. 2021; 433:166788

4. Laurino P, Tóth-Petróczy Á, Meana-Pañeda R, et al. An Ancient Fingerprint Indicates the Common Ancestry of Rossmann-Fold Enzymes Utilizing Different Ribose-Based Cofactors. PLOS Biol. 2016; 14:e1002396

5. Alva V, Söding J, Lupas AN. A vocabulary of ancient peptides at the origin of folded proteins. Elife 2015; 4:e09410

6. Sellés Vidal L, Kelly CL, Mordaka PM, et al. Review of NAD(P)H-dependent oxidoreductases: Properties, engineering and application. Biochim. Biophys. acta. Proteins proteomics 2018; 1866:327–347

7. Struck A-W, Thompson ML, Wong LS, et al. S-adenosyl-methionine-dependent methyltransferases: highly versatile enzymes in biocatalysis, biosynthesis and other biotechnological applications. Chembiochem 2012; 13:2642–2655

8. Kozbial PZ, Mushegian AR. Natural history of S-adenosylmethionine-binding proteins. BMC Struct. Biol. 2005; 5:19

9. Bastian S, Liu X, Meyerowitz JT, et al. Engineered ketol-acid reductoisomerase and alcohol dehydrogenase enable anaerobic 2-methylpropan-1-ol production at theoretical yield in Escherichia coli. Metab. Eng. 2011; 13:345–352

10. Hasegawa S, Uematsu K, Natsuma Y, et al. Improvement of the redox balance increases L-valine production by Corynebacterium glutamicum under oxygen deprivation conditions. Appl. Environ. Microbiol. 2012; 78:865–875

11. Scrutton NS, Berry A, Perham RN. Redesign of the coenzyme specificity of a dehydrogenase by protein engineering. Nature 1990; 343:38–43

12. Chánique AM, Parra LP. Protein Engineering for Nicotinamide Coenzyme Specificity in Oxidoreductases: Attempts and Challenges. Front. Microbiol. 2018; 9:194

13. Andreadeli A, Platis D, Tishkov V, et al. Structure-guided alteration of coenzyme specificity of formate dehydrogenase by saturation mutagenesis to enable efficient utilization of NADP+. FEBS J. 2008; 275:3859–69

14. Woodyer R, van der Donk WA, Zhao H. Relaxing the nicotinamide cofactor specificity of phosphite dehydrogenase by rational design. Biochemistry 2003; 42:11604–11614

15. Zheng H, Bertwistle D, Sanders DAR, et al. Converting NAD-specific inositol dehydrogenase to an efficient NADP-selective catalyst, with a surprising twist. Biochemistry 2013; 52:5876–83

16. Kallberg Y, Persson B. Prediction of coenzyme specificity in dehydrogenases/reductases. A hidden Markov model-based method and its application on complete genomes. FEBS J. 2006; 273:1177–1184

17. Cui D, Zhang L, Jiang S, et al. A computational strategy for altering an enzyme in its cofactor preference to NAD(H) and/or NADP(H). FEBS J. 2015; 282:2339–51

18. Cahn JKB, Werlang CA, Baumschlager A, et al. A General Tool for Engineering the NAD/NADP Cofactor Preference of Oxidoreductases. ACS Synth. Biol. 2017; 6:326–333

19. Gainza P, Sverrisson F, Monti F, et al. Deciphering interaction fingerprints from protein molecular surfaces using geometric deep learning. Nat. Methods 2020; 17:184–192

20. Geertz-Hansen HM, Blom N, Feist AM, et al. Cofactory: sequence-based prediction of cofactor specificity of Rossmann folds. Proteins 2014; 82:1819–28

21. Cheng H, Schaeffer RD, Liao Y, et al. ECOD: an evolutionary classification of protein domains. PLoS Comput. Biol. 2014; 10:e1003926

22. Sundararajan M, Taly A, Yan Q, et al. Rapid search for tertiary fragments reveals protein sequence-structure relationships. Protein Sci. 2015; 24:508–24

23. Ireland SM, Martin ACR. atomium-a Python structure parser. Bioinformatics 2020; 36:2750–2754

24. Salentin S, Schreiber S, Haupt VJ, et al. PLIP: fully automated protein-ligand interaction profiler. Nucleic Acids Res. 2015; 43:W443–7

25. Steinegger M, Söding J. MMseqs2 enables sensitive protein sequence searching for the analysis of massive data sets. Nat. Biotechnol. 2017; 35:1026–1028

26. Heinzinger M, Elnaggar A, Wang Y, et al. Modeling aspects of the language of life through transfer-learning protein sequences. BMC Bioinformatics 2019; 20:723

27. Kingma DP, Ba J. Adam: A Method for Stochastic Optimization. 2017;

28. Sundararajan M, Taly A, Yan Q. Axiomatic Attribution for Deep Networks. arXiv [cs.LG] 2017;

29. Schymkowitz J, Borg J, Stricher F, et al. The FoldX web server: an online force field. Nucleic Acids Res. 2005; 33:W382–8

30. Wang M, Zheng D, Ye Z, et al. Deep Graph Library: A Graph-Centric, Highly-Performant Package for Graph Neural Networks. 2019;

31. Lin T-Y, Goyal P, Girshick R, et al. Focal Loss for Dense Object Detection. IEEE Trans. Pattern Anal. Mach. Intell. 2020; 42:318–327

32. McInnes L, Healy J, Melville J. UMAP: Uniform Manifold Approximation and Projection for Dimension Reduction. 2018;

33. Shegay M V, Suplatov DA, Popova NN, et al. parMATT: parallel multiple alignment of protein 3D-structures with translations and twists for distributed-memory systems. Bioinformatics 2019; 35:4456–4458

34. Sali A, Blundell TL. Comparative protein modelling by satisfaction of spatial restraints. J. Mol. Biol. 1993; 234:779–815

35. Takase R, Mikami B, Kawai S, et al. Structure-based conversion of the coenzyme requirement of a short-chain dehydrogenase/reductase involved in bacterial alginate metabolism. J. Biol. Chem. 2014; 289:33198–33214

36. Nishiyama M, Birktoft JJ, Beppu T. Alteration of coenzyme specificity of malate dehydrogenase from Thermus flavus by site-directed mutagenesis. J. Biol. Chem. 1993; 268:4656–4660

37. Brinkmann-Chen S, Flock T, Cahn JKB, et al. General approach to reversing ketol-acid reductoisomerase cofactor dependence from NADPH to NADH. Proc. Natl. Acad. Sci. U. S. A. 2013; 110:10946–51

38. Chandrasekhar K, Wang J, Arjunan P, et al. Insight to the interaction of the dihydrolipoamide acetyltransferase (E2) core with the peripheral components in the Escherichia coli pyruvate dehydrogenase complex via multifaceted structural approaches. J. Biol. Chem. 2013; 288:15402–15417

39. Bocanegra JA, Scrutton NS, Perham RN. Creation of an NADP-dependent pyruvate dehydrogenase multienzyme complex by protein engineering. Biochemistry 1993; 32:2737–40

40. Petschacher B, Staunig N, Müller M, et al. Cofactor Specificity Engineering of Streptococcus mutans NADH Oxidase 2 for NAD(P)(+) Regeneration in Biocatalytic Oxidations. Comput. Struct. Biotechnol. J. 2014; 9:e201402005

41. Jensen CN, Ali ST, Allen MJ, et al. Mutations of an NAD(P)H-dependent flavoprotein monooxygenase that influence cofactor promiscuity and enantioselectivity. FEBS Open Bio 2013; 3:473–8

42. Bae B, Sullivan RP, Zhao H, et al. Structure and engineering of L-arabinitol 4-dehydrogenase from Neurospora crassa. J. Mol. Biol. 2010; 402:230–240

43. Chouhan BPS, Maimaiti S, Gade M, et al. Rossmann-Fold Methyltransferases: Taking a ‘β-Turn’ around Their Cofactor, S-Adenosylmethionine. Biochemistry 2019; 58:166–170

