## Supplementary Data for "Rossmann-toolbox: a deep learning-based protocol for the prediction and design of cofactor specificity in Rossmann-fold proteins"

Supplementary Methods, Figures and Tables

### Supplementary Methods

#### Data set augmentation

While training the sequence-based predictor, we used a data augmentation procedure, the purpose of which was to increase the model diversity and improve the calibration. In total we trained 250 models, using the procedure described in the main text (“*Prediction models”* in the Methods) and randomly varying the following parameters – a) maximum sequence identity in the training set (from 0.3 to 0.9 in 0.1 intervals), b) sequence coverage used to calculate sequence identity (from 0.5 to 0.8 in 0.1 intervals), c) amount of the noise applied to the labels (from 0 to 0.2 in 0.05 intervals, where the numbers denote the standard deviation of the zero-centered normal distribution from which the random noise samples were drawn from; individual labels were clipped to the original [0, 1] range after this operation). Additionally, we considered various batch sizes ranging from 8 to 40 in 8 intervals. The top 10 models, exhibiting the highest macro-F1 scores on the validation set, were used to create the final ensemble.

#### Development of the structure-based predictor

The limitation of the current Graph Neural Networks (GNNs) architectures in processing molecular data is their inability to jointly process information stored in nodes and edges [1]. This feature is essential for obtaining a complete graph representation of a protein structure in which residues (nodes) and interactions (edges) aren’t artificially separated and contain all the relevant information. In Graph Convolution Networks (GCNs) edges are used only to determine the radius of information flow. In the case of representing protein structures, this causes the loss of sequential information which is replaced with local structural information (residues connected according to their spatial proximity). Graph Attention Networks (GATs) are an extension of GCNs [2] in which edges not only define the local interactions but are also used to store temporary attention scores, known as edge weights, that are used to scale the importance of each connection.

In the task of representing the Rossmann βαβ motifs for use in GNN training, we are dealing with highly similar structures, whose corresponding graphs are frequently characterized by a similar topology (Figure 2 in the main text). Given this observation, we assumed that the features essential for the prediction of βαβ motifs interaction specificity may be defined by the nature of inter-residue interactions. Since such data cannot be captured by GCN and GAT architectures, we decided to expand the GAT algorithm with the possibility of handling regular edge features. As mentioned above, the original GAT idea is to compute individual attention scores, for each connection in the graph. The first step is to update node features with regular fully connected layer ${h^{'}}_{i}=Wh_{i} + b$, where $W$ and $b$ are learnable parameters, and to calculate attention scores using

$\boldsymbol{\alpha}_{\boldsymbol{ij}}\boldsymbol{= Softmax(}\boldsymbol{\epsilon}_{\boldsymbol{ij}}\boldsymbol{) =}\frac{\boldsymbol{exp(}\boldsymbol{\epsilon}_{\boldsymbol{ij}}\boldsymbol{)}}{\sum_{\boldsymbol{k\in N(i)}} \boldsymbol{.}\boldsymbol{exp(}\boldsymbol{\epsilon}_{\boldsymbol{ik}}\boldsymbol{)}}$	Eq. 1

$\boldsymbol{\epsilon}_{\boldsymbol{ij}}\boldsymbol{= LeakyReLU(A[}{\boldsymbol{h}^{\boldsymbol{'}}}_{\boldsymbol{i}}\boldsymbol{||}{\boldsymbol{h}^{\boldsymbol{'}}}_{\boldsymbol{j}}\boldsymbol{])}$	Eq. 2

where $a_{ij}$ denotes attention score, that is normalized weight over connection $n_{i}-n_{j}$, $\epsilon_{ij}$is the unnormalized weight where $A$ is a learnable matrix, $LeakyReLU$ is an activation function, and $(.||.)$ denotes vector concatenation operation. After obtaining the attention of each connection, updated feature vector ${h^{''}}_{i}$ of node $n_{i}$ is calculated with the equation

${\boldsymbol{h}^{\boldsymbol{''}}}_{\boldsymbol{i}}\boldsymbol{=}\sum_{\boldsymbol{k\in N(i)}} \boldsymbol{.}\boldsymbol{\alpha}_{\boldsymbol{ik}}{\boldsymbol{h}^{\boldsymbol{'}}}_{\boldsymbol{k}}$	Eq. 3

In this way, the individual GAT layers will learn which connections are important with respect to the information stored in $n_{i}$ and its neighbors. However, in such an implementation, the edge feature is only temporary and is not propagated further through the network. Our extension (Supplementary Figure 2) replaces equation (2) with

$\boldsymbol{\epsilon}_{\boldsymbol{ij}}\boldsymbol{= F}{\boldsymbol{f}^{\boldsymbol{'}}}_{\boldsymbol{ij}}$	Eq. 4

where $F$ is learnable matrix and ${f^{'}}_{ij}$ are updated features of edges produced by

${\boldsymbol{f}^{\boldsymbol{'}}}_{\boldsymbol{ij}}\boldsymbol{= LeakyReLU(A[}{\boldsymbol{h}^{\boldsymbol{'}}}_{\boldsymbol{i}}\boldsymbol{||}\boldsymbol{f}_{\boldsymbol{ij}}\boldsymbol{||}{\boldsymbol{h}^{\boldsymbol{'}}}_{\boldsymbol{j}}\boldsymbol{])}$	Eq. 5

in this case, the concatenation $(.||.||.)$ is performed over three quantities ${h^{'}}_{i}$, ${h^{'}}_{j}$, and $f_{ij}$. We decided to not use the resulting output edge features in the last EdgeGAT layer (in this case Eq. 4 is replaced with $\epsilon_{ij}={f^{'}}_{ij}$). The flow of the calculations is presented in Supplementary Figure 2 and can be summarized in four steps:

I. calculate auxiliary node feature vector ${h^{'}}_{i}$ (Supplementary Figure 2A)

II. use node (${h^{'}}_{i}$and ${h^{'}}_{j}$) and edge features ($f_{ij}$) for each existing connection in a graph to calculate output edge features ${f^{'}}_{ij}$ (Supplementary Figure 2B)

III. use output edge features ${f^{'}}_{ij}$ to calculate each edge attention $\alpha_{ij}$/$\epsilon_{ij}$ – connection importance factor. Note that importance coefficients are also calculated for self-loop edges connecting nodes with themselves (for clarity, these are not shown in Supplementary Figure 2C).

IV. sum node features ${h^{'}}_{i}$ multiplied by their importance factor to obtain output node features ${h^{''}}_{i}$. Owing to the usage of self-loops (see above), in cases when all the surrounding connections are irrelevant (attention score equals zero) then the new state ${h^{''}}_{i}$ will be the same as the old one ${h^{'}}_{i}$ (Supplementary Figure 2D).

#### Benchmark with alternative models

To verify whether the proposed EGAT architecture used for the **development of the structure-based predictor** provides better results than related GNN architectures, and to check how various parameters influence its efficiency, we performed a grid search analysis with the following parameters:

- type of GNN [GCN, GAT, and EGAT]
- number of GNN blocks [2, 3, and 4]
- size of node features in each block [32, 64, 128, 256, and 512]
- Cɑ distance threshold for defining the edges [6 Å to 13 Å]
- cross-validation (CV) fold [1 or 2] – the two CV folds were defined by swapping the test and validation sets

The 720 models, corresponding to all possible combinations of the aforementioned parameters, were trained with L2 regularization, varying learning rate, and stopping criteria depending on the macro-F1 score on the validation set, as described in the “*Prediction models”* section in the Methods*.* For each model, a macro-F1 score was calculated using the test set, and results were averaged over the two CV folds. As can be seen in Supplementary Figure 3, the EGAT architecture performed the best regardless of the Cɑ distance threshold. It is also noticeable that the superiority of EGAT over GCN and GAT grows along with the increase of the threshold, suggesting that edge features play a significant role in determining the importance of the individual edges (the proximity of Cɑ atoms does not imply that the residues they originate from form a significant interaction; however, such a significance can be derived from FoldX descriptors).

In selecting the Cɑ distance threshold for the calculation of the final models, we considered a trade-off between accuracy and computational time. On the one hand, the best EGAT model was obtained at the threshold of 10 Å; on the other, it was only slightly better (~1 % of macro-F1 score) than the second-best EGAT model obtained at 7 Å. This small gain in accuracy came at the cost of a considerable increase in the computational time required to train and run the model. This cost resulted from the fact that increasing the contact cut-off causes an increase in the number of graph edges (Supplementary Figure 4), and so, changing the cutoff from 7 Å to 10 Å is accompanied by almost the doubling the average number of edges.

To validate the effectiveness of combining the SeqVec embeddings with a deep neural network architecture for the **development of sequence-based predictor**, we trained complementary baseline models basing on the support-vector machines (SVM), K-nearest neighbors (kNN), and Random Forest (RFC) classifiers as implemented in *scikit-learn* Python package [3]. Models were trained on the SeqVec embedding as input and using the same dataset split as in the case of the deep learning-based predictor (Supplementary Table 2). Macro-F1 score on the validation set was used to determine optimal hyperparameters of the respective models and to select the best model, which was evaluated on the test set (Supplementary Figure 5). The obtained results indicated the poor performance of kNN and RFC classifiers (global macro-F1 scores of 0.78 and 0.81, respectively), and relatively good performance of the SVM classifier (global macro-F1 scores of 0.91). However, it must be emphasized that the performance of the SVM model is lower than that of the deep learning-based predictors (Figure 3 in the main text), especially in the case of distinguishing NAD and NADP binders (F1 scores shown in red in Figure 3 and Supplementary Figure 5).

#### Iterative mutational scan

For the identification of the complex mutations, that are composed of more than one mutation, an iterative mutational scan was performed employing the sequence-based method and Monte Carlo (MC) heuristics. In the implemented approach, the state of the system is fully described by the N point mutations (substitutions) applied to the WT sequence. In a single simulation step from 1 to N mutations are randomized (by changing their positions and substituting amino acids), and a new resulting sequence is evaluated with the sequence-based prediction model. A new state of the system is accepted according to the Metropolis criterium with the probability calculated with the following formula

$\boldsymbol{probability = min(1, exp((}\boldsymbol{S}_{\boldsymbol{N}}\boldsymbol{-}\boldsymbol{S}_{\boldsymbol{B}}\boldsymbol{)}\mathbf{/ kT))}$	Eq. 6

where $S_{N}$is the score of a sequence obtained in the given simulation step and $S_{B}$ is the score of the sequence obtained with the set of the currently best-performing mutations.

The convergence of such computations can be further enhanced with the use of modified probability distributions during the randomization procedure. In this case, the positions in the sequence are chosen according to the per-residue contributions to the prediction of the target cofactor, whereas the choice of amino acid in a given position is weighted based on position-specific scoring matrix (PSSM) scores derived from multiple sequence alignment (MSA) of βαβ motifs recognizing the desired cofactor (see also “*Per-residue contribution scores*” section in the main text). For each of the 38 WT βαβ motif sequences from the auxiliary test set (Supplementary Table 4), 50 independent simulations were performed with the following parameters: 500 MC steps, the N values ranging from 1 to 5, kT=0.05, and enabled enhanced probability distributions.

### Supplementary Figures

**Supplementary Figure 1.** The relationship between sequence-based and structure-based predictions for 1,647 cores from the train-validation-test set.

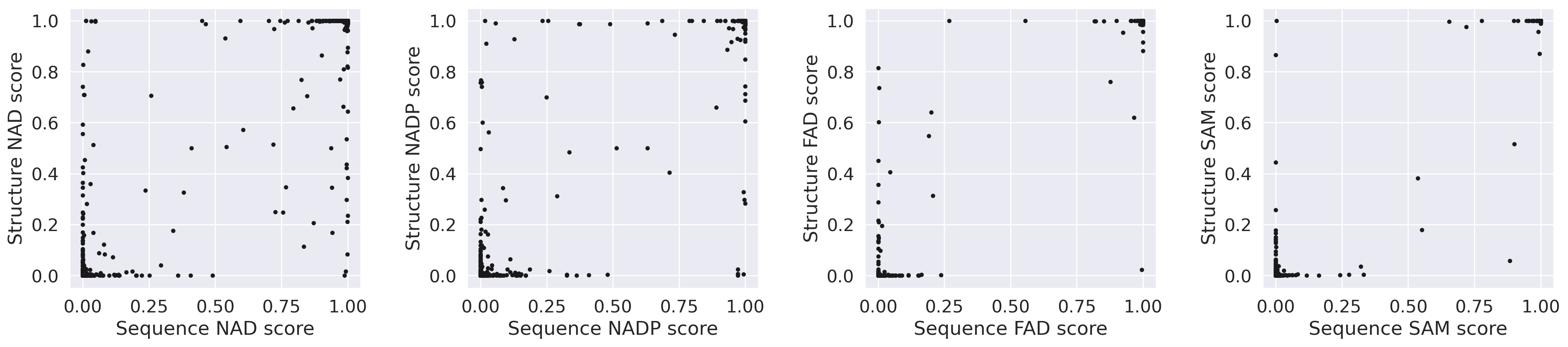

**Supplementary Figure 2.** Schema of a single Edge-GAT layer. For the sake of clarity, only some nodes and edges are annotated. **(A)** An exemplary input network comprising four nodes and three edges. h′_1_ and h′_2_ denote updated features (with regular fully connected) of nodes 1 and 2, respectively, whereas f_12_ denotes features of the edge that connects them. **(B)** Concatenation of node and edge features and calculation of updated edge features (f′_12_). **(C)** The updated edge features are used to calculate the importance (a value between 0 to 1) of node-edge-node connections (a′_12_). A new node feature (h′′_2_) is calculated as an average of surrounding node features weighted by the importance factors (a′). Note that the h′′_2_ is also used; its weighting importance factor is calculated from a self-looped edge feature (f′_22_). The self-loops were omitted for clarity. **(D)** Final output network with updated node and edge features.

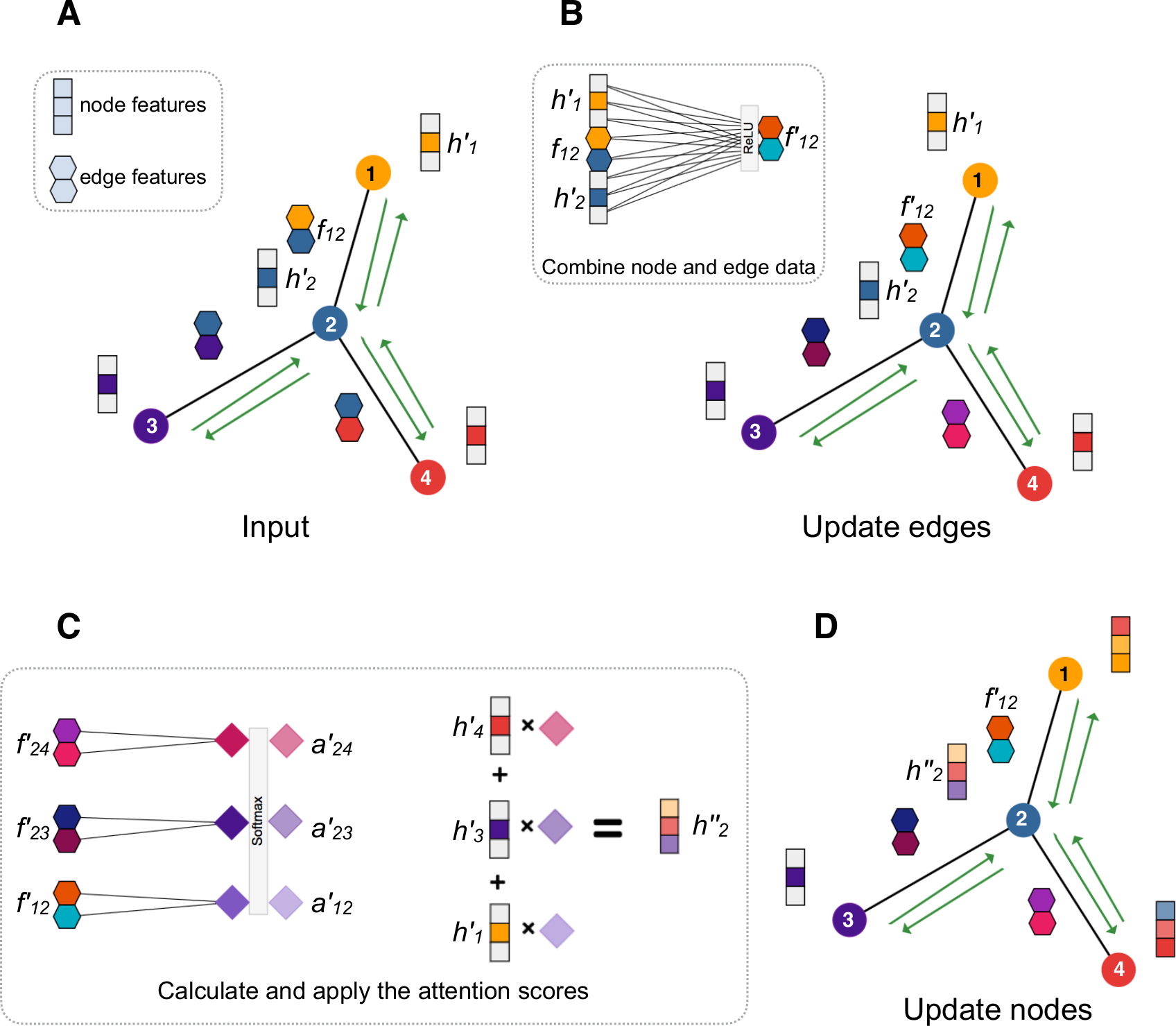

**Supplementary Figure 3.** Performance of GNN architectures (GCN, GAT, EGAT) for various cut-off used to define graph edges. The F1 scores were calculated using an independent test set.

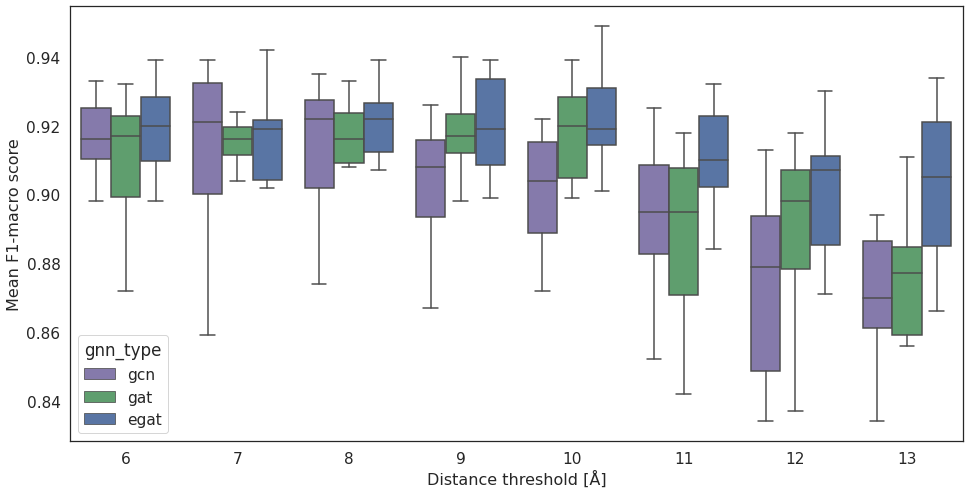

**Supplementary Figure 4.** Relationship between the cut-off used to define graph edges and their average number.

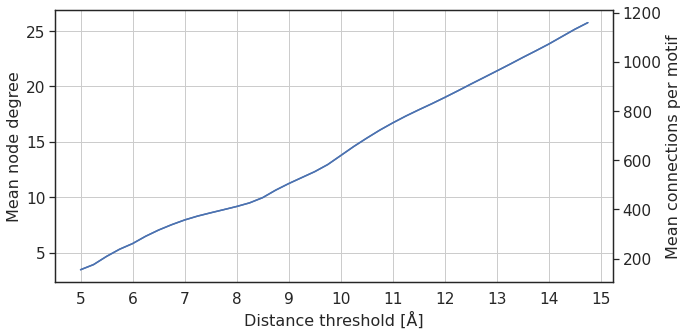

**Supplementary Figure 5.** Evaluation of alternative sequence-based prediction models using an independent test set. Each confusion matrix corresponds to a single method. The vertical and horizontal axes denote ground truth and predicted binding, respectively.

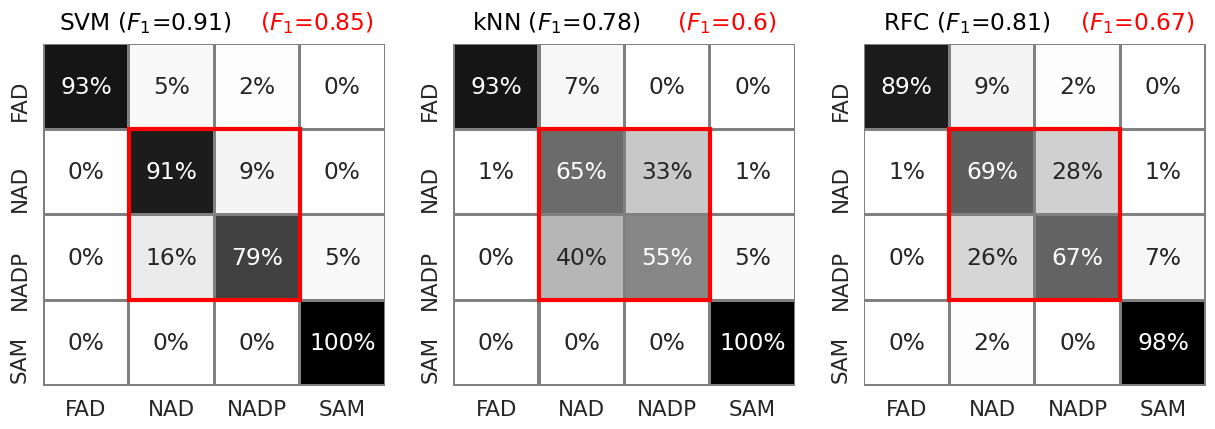

### Supplementary Tables

**Supplementary Table 1.** Query structures used to identify Rossmann fold structures in Protein Data Bank using MASTER [4]. Numbers in brackets define the residue ranges corresponding to β1, ɑ1, β2, β3, β4, and β5.

| **PDB id** | **Residue ranges** |
| --- | --- |
| 1xcb_A | (81, 86) (89, 96) (105, 111) (127, 128) (143, 146) (167, 170) |
| 1sqf_A | (250, 253) (258, 268) (272, 276) (298, 300) (318, 321) (369, 372) |
| 1bsv_A | (4, 9) (14, 24) (29, 32) (34, 35) (58, 61) (101, 105) |
| 2bh2_A | (289, 293) (300, 306) (310, 315) (336, 340) (359, 362) (383, 388) |
| 2bs2_A | (8, 11) (14, 26) (31, 34) (174, 176) (211, 214) (388, 390) |
| 1v9l_A | (212, 216) (219, 231) (235, 240) (245, 247) (294, 297) (316, 318) |
| 5cve_A | (64, 68) (73, 79) (86, 91) (111, 116) (129, 135) (160, 168) |
| 5i9n_A | (8, 11) (19, 31) (34, 39) (60, 62) (86, 90) (139, 144) |
| 3w6u_A | (3, 6) (11, 22) (25, 29) (45, 46) (59, 62) (89, 92) |
| 1inl_A | (94, 97) (102, 109) (116, 120) (147, 149) (166, 169) (200, 203) |
| 3fwz_A | (420, 423) (427, 439) (443, 447) (462, 465) (484, 487) (510, 513) |
| 4gmg_C | (8, 13) (18, 24) (32, 38) (56, 57) (69, 72) (96, 100) |
| 3fuu_A | (50, 53) (58, 67) (71, 75) (93, 96) (112, 116) (136, 141) |
| 3cin_A | (2, 7) (10, 25) (46, 53) (84, 85) (126, 129) (177, 181) |
| 1djq_A | (391, 395) (398, 411) (414, 418) (462, 464) (482, 485) (668, 671) |
| 1cjc_A | (8, 12) (16, 28) (34, 38) (74, 77) (97, 100) (362, 364) |
| 2nac_A | (192, 198) (200, 214) (216, 219) (236, 238) (249, 254) (278, 281) |
| 3pzr_A | (2, 6) (10, 22) (30, 35) (54, 55) (67, 70) (92, 95) |
| 3lcu_A | (129, 131) (136, 140) (152, 156) (176, 180) (194, 197) (223, 229) |
| 1pow_A | (216, 220) (226, 238) (240, 244) (257, 260) (278, 284) (301, 306) |
| 1bmd_A | (5, 9) (14, 24) (36, 39) (66, 69) (82, 84) (124, 126) |
| 2jfg_A | (8, 11) (14, 27) (32, 35) (52, 54) (67, 70) (90, 92) |
| 2hjr_A | (16, 20) (23, 36) (40, 44) (71, 74) (85, 88) (126, 129) |
| 1jw9_B | (34, 36) (45, 54) (58, 60) (103, 105) (124, 127) (149, 152) |
| 1oth_A | (190, 194) (199, 205) (214, 218) (244, 247) (259, 262) (299, 302) |
| 1dia_A | (167, 171) (177, 187) (191, 195) (198, 200) (211, 214) (232, 235) |
| 3anx_A | (80, 85) (89, 97) (103, 107) (134, 137) (152, 157) (189, 193) |
| 2ipx_A | (163, 167) (172, 183) (187, 191) (211, 214) (231, 236) (259, 262) |
| 2o23_A | (12, 16) (21, 33) (36, 41) (58, 62) (87, 89) (148, 153) |
| 3g8a_A | (83, 87) (94, 102) (106, 111) (132, 136) (152, 158) (179, 183) |
| 1c0p_A | (1006, 1010) (1013, 1026) (1029, 1033) (1156, 1159) (1174, 1177) (1326, 1328) |
| 1emd_A | (2, 6) (10, 23) (29, 33) (54, 58) (72, 74) (113, 115) |
| 1eiz_A | (54, 57) (63, 74) (78, 83) (93, 97) (120, 123) (160, 163) |
| 2o05_A | (96, 100) (105, 112) (119, 123) (149, 151) (168, 173) (202, 206) |
| 1e5q_A | (6, 9) (14, 24) (28, 32) (50, 52) (71, 73) (94, 96) |
| 3e7f_A | (38, 42) (45, 64) (69, 73) (104, 105) (155, 158) (213, 218) |
| 2glx_A | (3, 7) (10, 24) (27, 32) (50, 52) (66, 69) (89, 92) |
| 2zwa_A | (108, 114) (120, 127) (140, 146) (190, 194) (217, 224) (248, 255) |
| 1q7e_A | (12, 14) (19, 31) (34, 38) (70, 72) (94, 97) (122, 125) |
| 3pt6_A | (1142, 1147) (1152, 1162) (1164, 1170) (1188, 1190) (1222, 1225) (1265, 1271) |
| 4fu0_B | (4, 11) (16, 31) (36, 43) (49, 52) (100, 103) (107, 122) |
| 4dlk_B | (10, 14) (17, 30) (33, 38) (59, 69) (72, 75) (83, 92) |
| 3r5x_A | (1, 7) (10, 28) (33, 39) (42, 50) (55, 58) (62, 78) |
| 1nw7_A | (38, 44) (59, 64) (96, 106) (245, 248) (254, 264) (266, 271) |

**Supplementary Table 2.** Statistics of the train, test, and validation sets. Note that maximal sequence identity between any pair of βαβ motifs originating from two different sets never exceeds 30%.

| **Class** | **Cofactor** | **Number of βαβ motifs** |
| --- | --- | --- |
| Train | FAD | 262 |
|  | NAD | 306 |
|  | NADP | 274 |
|  | SAM | 264 |
| Test | FAD | 55 |
|  | NAD | 95 |
|  | NADP | 58 |
|  | SAM | 57 |
| Validation | FAD | 58 |
|  | NAD | 99 |
|  | NADP | 62 |
|  | SAM | 57 |

**Supplementary Table 3.** Structural features of nodes and edges extracted using FoldX *SequenceDetail* and *PrintNetworks* commands. In the network forward pass, the sequence and secondary structure elements were converted to randomly initialized vectors of fixed sizes 21 and 11. Such embeddings change in the training process for the better representation of stored information and are dense (unlike one-hot encoding). Float-type features (except for the learnable embeddings) were standardized by removing the mean and scaling to unit variance.

| Name | Source | Location | Size | Type |
| --- | --- | --- | --- | --- |
| Phi | *SequenceDetail* | Node | 1 | float |
| Psi | *SequenceDetail* | Node | 1 | float |
| Total | *SequenceDetail* | Node | 1 | float |
| backHbond | *SequenceDetail* | Node | 1 | float |
| sideHbond | *SequenceDetail* | Node | 1 | float |
| energy_VdW | *SequenceDetail* | Node | 1 | float |
| Electro | *SequenceDetail* | Node | 1 | float |
| energy_SolvP | *SequenceDetail* | Node | 1 | float |
| energy_SolvH | *SequenceDetail* | Node | 1 | float |
| energy_vdwclash | *SequenceDetail* | Node | 1 | float |
| entrop_sc | *SequenceDetail* | Node | 1 | float |
| entrop_mc | *SequenceDetail* | Node | 1 | float |
| cis_bond | *SequenceDetail* | Node | 1 | float |
| energy_torsion | *SequenceDetail* | Node | 1 | float |
| backbone_vdwclash | *SequenceDetail* | Node | 1 | float |
| energy_dipole | SequenceDetail | Node | 1 | float |
| Sidechain Contact Ratio | SequenceDetail | Node | 1 | float |
| Mainchain Contact Ratio | SequenceDetail | Node | 1 | float |
| residue hydrophobicity [5] | own | Node | 1 | float |
| residue volume [6] | own | Node | 1 | float |
| residue embedding | own | Node | 21 | float |
| DSSP-based secondary structure embedding [7] | own | Node | 11 | float |
| Hydrogen bonds | *PrintNetworks* | Edge | 4 | binary |
| Volumetric | *PrintNetworks* | Edge | 4 | binary |
| Electro | *PrintNetworks* | Edge | 4 | binary |
| VdWClashes | *PrintNetworks* | Edge | 4 | binary |
| Cɑ–Cɑ inverse distance | own | Edge | 1 | float |
| side chain angle | own | Edge | 1 | float |
| is Cɑ–Cɑ contact structural | own | Edge | 1 | binary |
| is Cɑ–Cɑ contact sequential | own | Edge | 1 | binary |

**Supplementary Table 4.** Auxiliary test set comprising 38 experimentally confirmed cases of switching cofactor specificity between NAD and NADP by point (*p*) or complex (*c*) mutations. The D38Q NAD 🡪 NADP mutation of *Drosophila melanogaster* alcohol dehydrogenase was obtained from [8,9] where it was listed with a reference to [10] which appears to be wrong. For this reason, no source is provided in this case.

|  | Source | WT Cof. | MUT. Cof. | PDB_chain | Mutations | Type |
| --- | --- | --- | --- | --- | --- | --- |
| 1 | Bae et al., 2010 | NAD | NADP | 3m6i_A | D211S, I212R | *c* |
| 2 | Ehrensberger et al., 2006 | NAD | NADP | 1zem_A | D38S, M39R | *c* |
| 3 | Takase et al., 2014 | NAD | NADP | 4tkm_A | T16S, E17Q, N37H, S38G, H39R, V40K, D41A | *c* |
| 4 | Bastian et al., 2011 | NADP | NAD | 3ulk_A | S78D | *p* |
| 5 | Brinkmann-Chen et al., 2013 | NADP | NAD | 4tsk_A | R48P, S51L, S52D | *c* |
| 6 | Brinkmann-Chen et al., 2013 | NADP | NAD | 4kqw_A | S61D, S63D | *c* |
| 7 | Cahn et al., 2016 | NADP | NAD | 1piw_A | S210D, R211P, K215E | *c* |
| 8 | Nakanishi et al., 1997 | NADP | NAD | 1cyd_A | T38D | *p* |
| 9 | Rane et al., 1997 | NADP | NAD | 3ulk_A | R68D, K69L, K75V, R76D | *c* |
| 10 | Takase et al., 2014 | NADP | NAD | 4w7i_A | H37N, G38S, R39H, K40V, A41D | *c* |
| 11 | Zhang et al., 2009 | NADP | NAD | 3ctm_A | S67D, H68D | *c* |
| 12 | Andreadeli et al., 2008 | NAD | NADP | 2j6i_A | D195Q, Y196H | *c* |
| 13 | Wu et al., 2009 | NAD | NADP | 2j6i_A | D195Q, Y196R, Q197N | *c* |
| 14 | Carrigan et al., 2007 | NAD | NADP | 2yfq_A | E243K | *p* |
| 15 | Kumar et al., 2017 | NADP | NAD | 5n2i_A | S50E | *p* |
| 16 | Bocanegra et al., 1993 | NAD | NADP | 4jdr_A | E205V, M206R, F207K, D208H, P212R | *c* |
| 17 | Scrutton et al., 1990 | NADP | NAD | 1ger_A | A179G, A183G, V197E, R198M, K199F, H200D, R204P | *c* |
| 18 | Woodyer et al., 2003 | NAD | NADP | 4e5n_A | E175A, A176R | *c* |
| 19 | Cui et al., 2015 | NAD | NADP | 3awd_A | Q20R, D43S | *c* |
| 20 | Holmberg et al., 1999 | NAD | NADP | 1ldn_A | I51K, D52S | *c* |
| 21 | Bernard et al., 1995 | NAD | NADP | 1j49_A | D176A | *p* |
| 22 | Dambe et al., 2006 | NADP | NAD | 2glx_A | A13G, S33D | *c* |
| 23 | ? | NAD | NADP | 1mg5_A | D38Q | *p* |
| 24 | Cahn et al., 2016 | NADP | NAD | 3doj_A | R31L, T32K, K35D | *c* |
| 25 | Clermont et al., 1993 | NAD | NADP | 1gd1_O | D32N | *p* |
| 26 | Hasegawa et al., 2012 | NADP | NAD | 6jx2_A | S34G, L48E, R49F | *c* |
| 27 | Rosell et al., 2003 | NADP | NAD | 1p0c_A | G1222D, T1223I, H1224N | *c* |
| 28 | Kristan et al., 2007 | NADP | NAD | 3qwf_A | Y49D | *p* |
| 29 | Pick et al., 2014 | NADP | NAD | 1uuf_A | T205D, T206I, S207N | *c* |
| 30 | Schepens et al., 2000 | NADP | NAD | 1civ_A | G80D, S81I, R83Q, S84A | *c* |
| 31 | Bubner et al., 2008 | NAD | NADP | 1m2w_A | E68K, D69A | *c* |
| 32 | Capone et al., 2012 | NAD | NADP | 1bgv_A | F238S, P262S, D263K | *c* |
| 33 | Gul-Karaguler et al., 2001 | NAD | NADP | 2nad_A | D221S | *p* |
| 34 | Jensen et al., 2013 | NAD | NADP | 4a9w_A | Q193R, H194T | *c* |
| 35 | Nishiyama et al., 1993 | NAD | NADP | 1bmd_A | E41G, I42S, P43E, Q44R, A45S, M46F, K47Q | *c* |
| 36 | Petschacher et al., 2014 | NAD | NADP | 2bc0_A | D190A, V191R, V192H, A197R | *c* |
| 37 | Zheng et al., 2013 | NAD | NADP | 3nt2_A | A12K, D35S, V36R | c |
| 38 | García-Guevara et al., 2017 | NADP | NAD | 1nyt_A | N149D, V152F, S131A, L135A | c |

**Supplementary Table 5.** Results of iterative mutational scan for the auxiliary test set cases featuring two or more mutations. Numbers in brackets define the position of a given experimentally confirmed (“correct”) mutation/coupling in the ranking. The question mark sign indicates that none of the “correct” mutations/couplings were among the top 20 predictions.

|  | **Predicted mutations** | **Predicted couplings** |
| --- | --- | --- |
| 1 | I212R (1), D211S (2) | D211S-I212R (1) |
| 2 | M39R (1), D38S (2) | D38S-M39R (1) |
| 3 | H39R (1), V40K (3), S38G (8) | H39R-V40K (2), S38G-H39R (10) |
| 5 | R48P (5), S52D (15), S51L (18) | N/A |
| 6 | N/A | N/A |
| 7 | S210D (1), R211P (9) | S210D-R211P (8) |
| 9 | R68D (14) | N/A |
| 10 | K40V (11) | N/A |
| 11 | S67D (10) | N/A |
| 12 | N/A | N/A |
| 13 | Y196R (4) | N/A |
| 16 | E205V (17) | N/A |
| 17 | A179G (19) | N/A |
| 18 | A176R (6), E175A (10) | N/A |
| 19 | D43S (4) | N/A |
| 20 | D52S (2) | N/A |
| 22 | S33D (1), A13G (5) | A13G-S33D (11) |
| 24 | R31L (6) | N/A |
| 26 | L48E (2), R49F (6) | N/A |
| 27 | G1222D (1) | N/A |
| 29 | T205D (1), T206I (2) | T205D-T206I (1) |
| 30 | G80D (1), S81I (20) | N/A |
| 31 | E68K (7), D69A (10) | N/A |
| 32 | D263K (13), P262S (19) | N/A |
| 34 | Q193R (2) | N/A |
| 35 | E41G (12), I42S (13), Q44R (19) | N/A |
| 36 | D190A (4), V191R (15) | N/A |
| 37 | D35S (3), V36R (4) | D35S-V36R (1) |
| 38 | N149D (1) | D211S-I212R (1) |

### Bibliography

1. Gupta A, Matta P, Pant B. Graph neural network: Current state of Art, challenges and applications. Mater. Today Proc. 2021;

2. Veličković P, Cucurull G, Casanova A, et al. Graph Attention Networks. 2017;

3. Pedregosa F, Varoquaux G, Gramfort A, et al. Scikit-learn: Machine Learning in Python. J. Mach. Learn. Res. 2011; 12:2825–2830

4. Sundararajan M, Taly A, Yan Q, et al. Rapid search for tertiary fragments reveals protein sequence-structure relationships. Protein Sci. 2015; 24:508–24

5. Kyte J, Doolittle RF. A simple method for displaying the hydropathic character of a protein. J. Mol. Biol. 1982; 157:105–32

6. Zamyatnin AA. Protein volume in solution. Prog. Biophys. Mol. Biol. 1972; 24:107–23

7. Kabsch W, Sander C. Dictionary of protein secondary structure: pattern recognition of hydrogen-bonded and geometrical features. Biopolymers 1983; 22:2577–637

8. Cui D, Zhang L, Jiang S, et al. A computational strategy for altering an enzyme in its cofactor preference to NAD(H) and/or NADP(H). FEBS J. 2015; 282:2339–51

9. Marohnic CC, Bewley MC, Barber MJ. Engineering and characterization of a NADPH-utilizing cytochrome b5 reductase. Biochemistry 2003; 42:11170–82

10. Bocanegra JA, Scrutton NS, Perham RN. Creation of an NADP-dependent pyruvate dehydrogenase multienzyme complex by protein engineering. Biochemistry 1993; 32:2737–40
